# Yawning reveals energy-dynamics mismatch and neural inertia across state transitions

**DOI:** 10.64898/2026.04.26.720882

**Authors:** Xiangshang Meng, Qi Sun, Muzi Li, Xuanze Li, Guangtai Liang, Ling Qin, Wenfei Tan

## Abstract

Yawning is a highly conserved behavior, yet its neural dynamics across arousal state transitions remain poorly understood. Classical reflex models fail to account for the precise timing of yawns during large-scale changes in brain network organization. Here, we investigated the neural signatures of yawning using simultaneous electroencephalography (EEG), electromyography (EMG), and kinematic recordings in a beagle model during wakefulness and propofol anesthesia. Propofol-associated yawning exhibited marked temporal asymmetry, with 89.2% of events occurring during emergence from anesthesia. To resolve the fast neural dynamics surrounding these events, we applied empirical mode decomposition coupled with the Hilbert–Huang transform (EMD-HHT) analyses. Across induction, recovery, and wakefulness, yawning was consistently associated with a transient decoupling between local and global network dynamics: a brief increase in local γ-band power coincided with a drop in network flexibility, quantified by instantaneous frequency volatility; IFV). We termed this reproducible motif as “Energy-Dynamics Mismatch” (EDM). A machine learning classifier leveraging EDM features reliably decoded the brain state immediately preceding yawning. We further validated the presence of this signature in an independent human cohort during anesthesia induction. These findings indicate that yawning is not a passive reflex, but is tightly linked to moments of network instability during state transitions. From a computational perspective, EDG may reflect a transient “control effort” that preserves information integration in the presence of “neural inertia.” This dynamic signature may provide a quantitative biomarker for monitoring arousal transitions brain-computer interface (BCI) interface state-alerting.

**Significance Statement:** Yawning is a ubiquitous and evolutionarily conserved behavior, yet its neural basis remains elusive. Using high-resolution neural recordings in a cross-species model, we show that yawning is not a passive reflex, but reliably coincides with a distinct, transient brain-state signature. Specifically, yawning occurs when local high-frequency cortical activity briefly increases while global network flexibility decreases – a pattern we term an “Energy-Dynamics Mismatch” (EDM). We propose that EDM reflects a compensatory neural process engaged during moments of network instability, consistent with overcoming neural inertia to preserve information integration. By identifying this conserved neural marker, this work advances the biological understanding of yawning and suggests a potential biomarker for detecting critical brain states in future BCI.

## Introduction

Yawning is an evolutionarily conserved physiological behavior observed across vertebrate species (1, 2). It frequently occurs during transitions in brain state, particularly across the sleep-wake cycle (3) and changes in anesthesia depth (4). Classic theories have primarily attributed yawning to brain thermoregulation (5) or airway homeostasis reflexes (6). However, these accounts struggle to explain the remarkably precise timing of yawns during narrow windows of neuronal state transition.

Contemporary neuroimaging research predominantly focuses on activation patterns of macroscopic brain regions (7, 8). As a result, it lacks the temporal resolution necessary to resolve millisecond-scale neural processes that precede spontaneous physiological behaviors such as yawning. To overcome this, we employed a propofol-induced anesthesia–emergence paradigm, which offers a well-controlled and highly reproducible model of transient brain-state transitions and graded levels of consciousness. Propofol-induced state transitions exhibit an asymmetric resistance between induction and recovery, known as ‘neural inertia’ (9, 10). Despite its robust macroscopic characterization, it remains unclear how this macroscopic pharmacodynamic phenomenon maps onto microscopic brain network dynamics. In particular, whether the neuronal microstates preceding yawning reflect an active attempt by the brain to overcome this hysteretic resistance has not been empirically demonstrated.

Here, we hypothesize that yawning reflects latent neuronal computations that occur as the brain network negotiates transitions across different levels of consciousness. Cortical energy measures (e.g., γ power) (11) and network dynamic metrics (e.g., instantaneous frequency volatility, IFV) (12) are typically studied in isolation. Their potential interaction during brief, transitional time periods has received little attention. Neural variability theory posits that the degrees of freedom of a neural network are partially captured by dynamic fluctuations in neural signals, reflecting the network’s capacity to explore and switch between functional states (13, 14). IFV offers a quantitative measure of this process by indexing the moment-to-moment reorganization of oscillatory frequency (15, 16). Accordingly, we define IFV as a metric of “microscale neuronal variability,” indirectly reflecting the intrinsic flexibility of local network state transitions.

By integrating cordical energy and network flexibility within a unified observational framework, we identified a significant dissociation between these two elements in the microscopic time window preceding yawning across multiple arousal states. Specifically, cortical energy increased while network flexibility simultaneously decreased, indicating a transient internal conflict within the system. We term this phenomenon “Energy-Dynamics Mismatch” (EDM). This mismatch can be conceptualized as a form of neural “control effort” deployed in response to macro-state conflicts (17, 18).

To test this theory, we developed a standardized anesthetic state transition model in beagle dogs and simultaneously recorded multimodal neurophysiological and high-precision kinematic signals. Compared with rodents, beagles possess more developed cortical architecture and network dynamics that more closely approximate those of humans. We observed a pronounced temporal asymmetry in propofol-induced yawning, with strong clustering during the anesthesia recovery period. Using high-resolution nonlinear time–frequency analysis, we characterized the underlying microscopic dynamics. Across all conditions, including deep anesthesia, delayed recovery, and normal wakefulness, yawning was consistently preceded by an increase in local cortical γ power alongside a reduction in network flexibility, as measured by instantaneous frequency volatility (IFV). These findings indicate that EDM represents a universal microstate feature preceding yawning. Notably, this mismatch persisted even when the specific intrinsic mode function (IMF) associated with reduced flexibility shifted its center frequency to align with global brain rhythms, such as anesthesia-induced slowing (19). We subsequently verified this dynamic decoupling pattern in an independent human cohort, demonstrating cross-species generalizability.

Together, these findings redefine the neural precursor states associated with yawning. This process provides an objective, high-resolution quantitative framework for understanding the contextual and computational underpinnings of this evolutionarily conserved behavior. Beyond mechanistic insight, EDM may serve as a novel biomarker of transitional brain states and offers a potential foundation for future applications, including BCI early warning systems based on microscopic dynamic tension.

## Results

### Propofol-Induced Yawning Exhibits Dose-Dependence and Recovery-Phase Specificity

We first conducted preliminary dose-determining experiments to explore the temporal distribution and neurodynamic characteristics of yawning during anesthetic transitions (Fig. 1A). We contrasted a suggested induction dose (8 mg/kg), delivered by intravenous infusion over one minute, with a half-dose (4 mg/kg). To ensure spatial accuracy during neuroelectrophysiological recordings, we developed an individualized MRI-guided stereotactic targeting workflow. Titanium screw artifacts at the bregma served as a physical reference point. Using high-precision 3D brain reconstruction, we precisely mapped the spatial coordinates of four major brain regions (Fig. S1A), providing reliable epidural spatial anchors for subsequent recordings.

**Figure 1.**
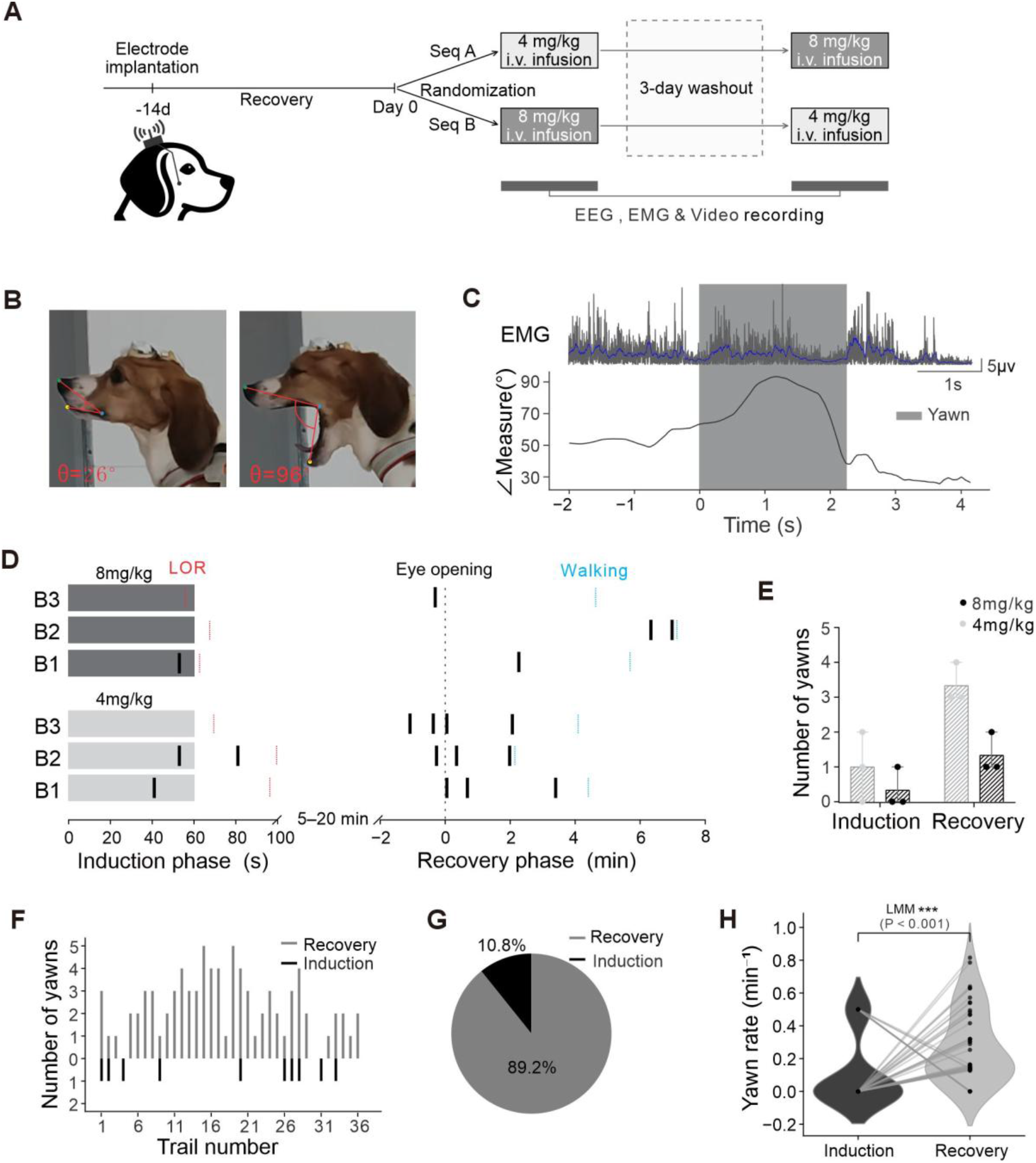
Kinetic definition of propofol-induced yawning and temporal asymmetry during state transitions. (A) A schematic showing the plan for the first experiment to find the right dose (n = 3). (B) How we defined a yawn using DeepLabCut to track points on the face. This shows the change in mouth shape from closed (θ = 26°) to the widest point (θ = 96°) in one typical beagle. (C) Matching the behavior and electrical signals in time. The mouth-opening angle (black line), the raw masseter electromyography (EMG, grey), and the blue curve are synchronized in the time domain. Grey boxes show yawns that lasted longer than 1 s, where t = 0 is the start of the action. (D) Plots showing when yawns (black ticks) happened at different doses. Dashed vertical lines show the loss of response (LOR, red), when eyes opened (black), and when walking started again (blue). (E) Number of yawns during the induction and recovery for two doses. Note: These data show a general trend only. (F) A mirror plot for the formal experiment (4 mg/kg, 36 trials) showing the distribution of yawns during recovery (top, grey) and induction (bottom, black). (G) The percentage of yawns that happened in induction compared to recovery. (H) Comparing yawn rates after adjusting for time. In the violin plot, grey lines connect the data from the same trial. ***P < 0.001, using a linear mixed-effects model (LMM) that treats each animal as a random factor.

We then developed a multimodal quantitative analysis pipeline combining neurophysiological (EEG/EMG) recordings with high-resolution kinematic measurements. The geometric characteristics of yawning were quantified through dynamic tracking of facial vectors using DeepLabCut (Fig. 1B). This mechanical facial movement was strictly time-locked to masseter EMG activities, as shown in Figure 1C. According to our definition, a valid yawn is a synergistic motor-electrophysiological event lasting longer than one second and accompanied by high-amplitude EMG bursts. Using this multimodal alignment strategy, the kinetic onset (t=0) of each yawning event could be specified objectively.

When yawn occurrences were projected onto the anesthetic timeline, a highly imbalanced temporal distribution with clear dose-dependent tendencies emerged. The induction phase, spanning drug administration to loss of response (LOR), showed a sparse activity with only occasional events in the raster plot (Fig. 1D). In contrast, dense, cluster-like bursts were observed during the recovery phase. Descriptive analysis indicated that the 4 mg/kg dose elicited more yawns than the 8 mg/kg dose (Fig. 1E). Based on these observations, we selected 4 mg/kg as the standard dose for all subsequent formal trials.

These initial findings were supported by quantitative data from 36 independent trials at the selected dose. The temporal discrepancy is illustrated in a mirror raster plot (Fig. 1F), in which dense gray vertical bars during recovery stand in sharp contrast to the extremely sparse black bars during induction. This distribution demonstrated clear time-window specificity: the majority of yawns occurred between eye-opening and walking. Overall, only 10.8% of documented yawns occurred during induction, compared to 89.2% during recovery (Fig. 1G).

To account for unequal phase durations and minimize exposure bias, absolute event counts were converted to time-normalized occurrence rates (events/min). For formal statistical inference, we applied a linear mixed-effects model (LMM) with individual-level random effects to account for the nested structure of repeated measurements. The time-corrected yawning rate was significantly higher during recovery than during induction (P < 0.001; Fig. 1H).

This statistically corrected temporal asymmetry quantitatively suggests the presence of neurodynamic hysteresis in yawning behavior. Notably, yawning preferentially coincided with the “rebooting” of consciousness networks rather than their “shutdown.” These dense recovery-phase events may reflect a dynamic interplay between rising arousal demands and residual “Neural Inertia” within the underlying network.

### Neural Oscillatory Reconfiguration and State-Space Trajectories in Propofol-Induced Anesthesia

We used a longitudinal, multi-cycle repeated-measures paradigm (n = 6; Fig. 2A). Following electrode implementation and recovery, each animal underwent three standardized anesthesia-emergence cycles, yielding 36 anesthesia recordings and 96 anesthesia-related yawning events. This design ensured strong statistical power while minimizing the neuropharmacological impact of drug accumulation.

**Figure 2.**
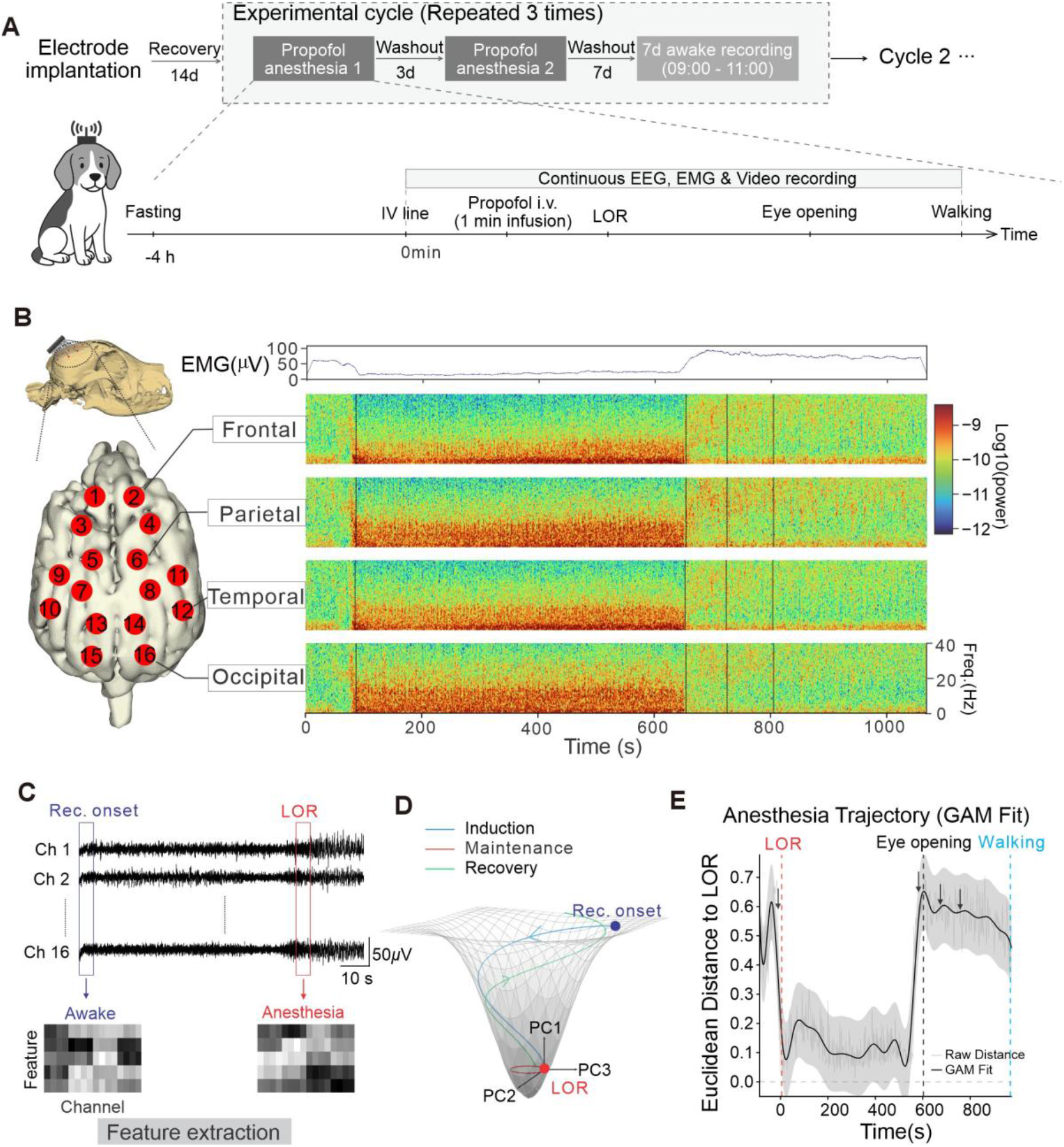
Analysis of neural state trajectory dynamics during propofol induction and recovery. (A) A drawing of the experiment and key points in behavior (n = 6). These points include loss of response (LOR), opening eyes, and starting to walk again. (B) Spatial electrode distribution and macro-scale time-frequency evolution. Left: A map showing 16 EEG channels across four main areas of the brain. Top right: A typical muscle signal (EMG) wave. Bottom right: Changes in the power spectrum for one channel from being awake, through anesthesia, and then recovery. Black dashed lines show when yawns happened. (C) Extraction process for neural state features. Blue and red boxes show the 5-second windows we used to find the features for being awake (Rec. onset) and for the loss of response (LOR). (D) 3D paths of brain states using Principal Component Analysis (PCA). The PC1, PC2, and PC3 axes show the first three components of the state subspace. Blue, red, and green lines show the induction, maintenance, and recovery phases of the experiment. (E) Euclidean distance dynamics between instantaneous neural states and the LOR centroid. Time zero on the x-axis is the time of LOR. Grey lines show the raw distance data. The black line and the shaded area show the Generalized Additive Model (GAM) fit and its 95% confidence interval (CI). Black arrows point to yawning events.

Intravenous propofol infusion rapidly induced anesthesia, as evidenced by a marked reduction in electromyography (EMG) amplitude (Fig. 2B, top right). In the time-frequency domain, global brain oscillatory patterns were substantially reconfigured (Fig. 2B, bottom right). Low-frequency δ (1–4 Hz) and α (8–13 Hz) power increased throughout induction (0–100 s), whereas high-frequency γ (30–40 Hz) power decreased. These changes were uniform across cortical regions, remained stable during the maintenance phase, and gradually returned to waking levels during recovery.

To better characterize the continuous evolution of brain states, we constructed a multidimensional neural state-space based on power spectral density across five canonical frequency bands. For trajectory quantification, wakefulness and loss of response (LOR) were defined as reference anchors (Fig. 2C). Principal component analysis (PCA) was used to project the original 80-dimensional feature space onto a 3-dimensional subspace, revealing highly structured state trajectories. Anesthetic states converged markedly into a compact region near the LOR centroid, whereas wakeful states were more topologically distributed (Fig. 2D). The empirical dataset provided topological support for both this “funnel-like” convergence and the nonlinear dynamics of state transitions (Fig. S1B). Notably, although the recovery trajectory (green) eventually returned to the wakefulness region, it remained spatially distinct from the induction path (blue). This reveals a pronounced state-space imbalance across the anesthesia-emergence process. Such hysteresis in network evolution trajectories provides a macro-electrophysiological representation of “neural inertia”.

To further quantify these transitions, we computed the Euclidean distance between the instantaneous neuronal state and the LOR centroid. The resulting distance time-series was subsequently fitted using generalized additive models (GAM) (Fig. 2E). Three key observations emerged from the GAM fits. First, during induction, the Euclidean distance rapidly decreased, closely coinciding with the behavioral onset of LOR. Second, during maintenance, the distance remained at a low, stable level, consistent with neurodynamics trapped in a deeply inhibited state. Third, upon recovery, the distance rose sharply at the moment of “eye-opening”. However, the model revealed that a high-level steady state between “eye-opening” and “walking” was not sustained. Instead, the neural state distance gradually declined during this interval (Fig. 2E, black arrows), suggesting that yawning may serve as an objective behavioral marker of underlying network reconfiguration when the brain encounters dynamic barriers during state transitions.

### Unsupervised Learning-Based Deconstruction of Yawning Neural Phenotypes and State-Manifold Mapping

To more finely dissect the micro-neural dynamics associated with yawning, we applied unsupervised spectral phenotypic classification. We extracted 64-dimensional feature vectors from 16 EEG channels, comprising global absolute power in the δ (1–4 Hz) and γ (30–45 Hz) bands, as well as their temporal changes before and after yawning. K-means clustering combined with UMAP dimensionality reduction revealed that the neurodynamic backgrounds of yawning segregated into four topologically distinct clusters (Clusters 0–3; Fig. 3A).

**Figure 3.**
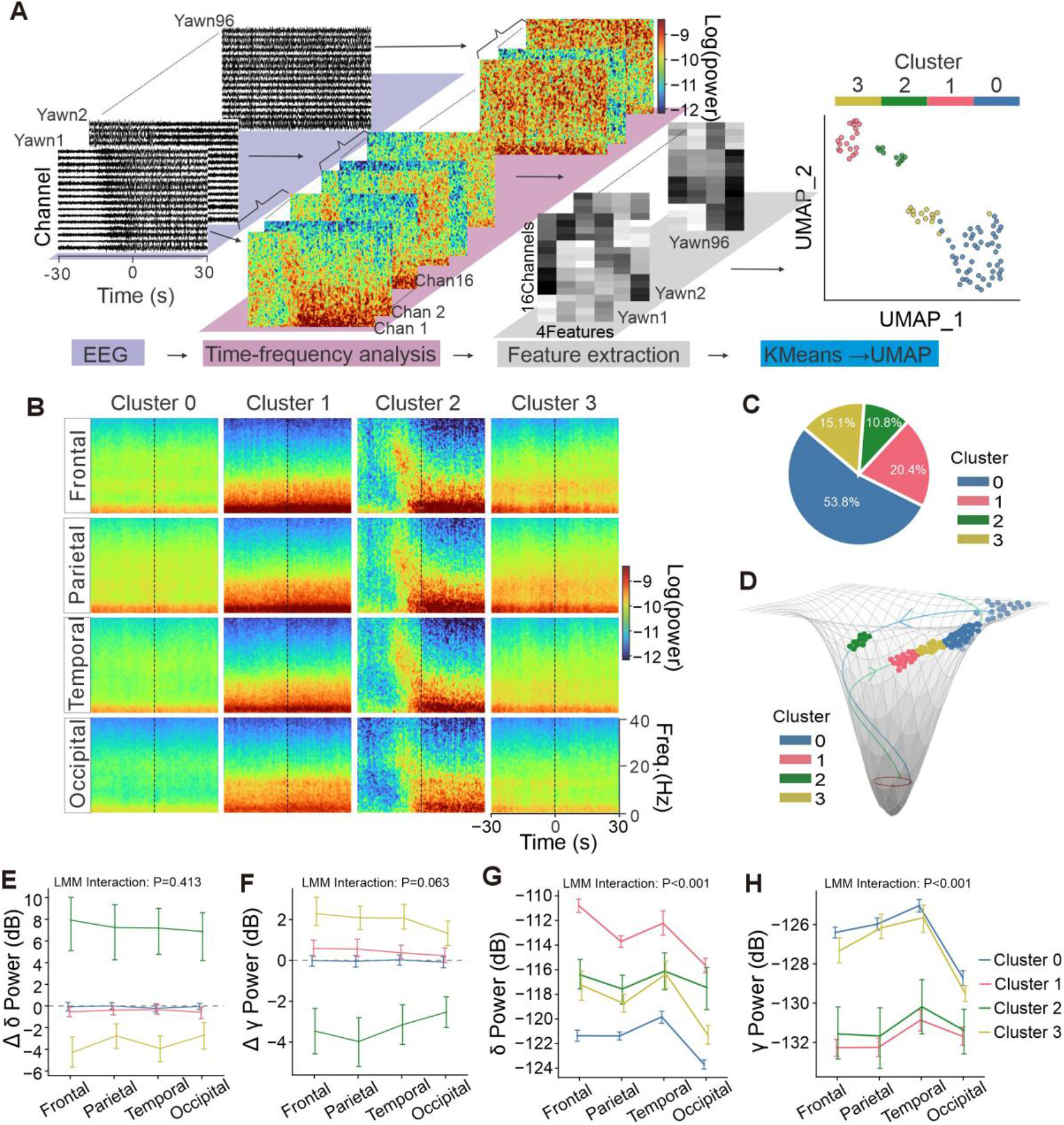
Classifying the neural spectral patterns of yawning and their mapping in state space. (A) The steps for unsupervised clustering and UMAP projection to find yawn types (Clusters 0–3). We calculated the features using the average power and the relative change (Δ) of the δ and γ bands in the time window around the yawn (± 30 s). (B) Average time-frequency maps (spectrograms) for the four clusters in four main brain areas. The vertical black dashed line marks the start of the yawning action (t = 0). (C) The portion of each cluster out of all the yawning events that were recorded. (D) Map of yawning events on a 3D brain state shape (manifold). The color of each dot shows which cluster it belongs to. The blue, red, and green lines represent the induction, maintenance, and recovery phases of the experiment. (E–H) Mean relative power changes (E, F) and total average power (G, H) for the δ and γ bands. The plots show the mean ± 95% CI for each cluster across the four brain regions. P-values show the “Cluster × Region” interaction effect calculated with a linear mixed-effects model (LMM).

Average spectrograms across frontal, parietal, temporal, and occipital cortical regions revealed distinct spectral evolution patterns for each cluster (Fig. 3B). Before and after yawning, Cluster 0, the most prevalent phenotype (53.8% of all events; Fig. 3C), consistently exhibited low δ power and high-frequency γ activity. This pattern suggests occurrence in a state approximating the wakeful baseline during late recovery. In contrast, Cluster 2 (10.8%) displayed a characteristic induction-phase collapse, marked by a simultaneous surge in δ power and a pronounced reduction in γ activity, indicative of a rapid descent into deep anesthesia. The opposite progression was observed in Cluster 3 (15.1%), which showed decreasing δ power alongside recovery of γ activity, consistent with early recovery dynamics. Cluster 1 (20.4%) was dominated by high δ and low γ power, reflecting the macro-electrophysiological signature of deep anesthesia. The occurrence of yawning under such pronounced suppression suggests a hidden micro-dynamic transition, implying that beneath an apparently stable inhibitory state, the cortex may undergo spontaneous reconfiguration.

Mapping these four clusters into the 3D neural state manifold revealed that yawning events closely follow δ and γ topological organization. Cluster 2 aligned with the descending induction trajectory, while Cluster 1 localized to the manifold base within a deeply metastable inhibited region. Along the ascending recovery trajectory, Clusters 0 and 3 progressed upward toward wakefulness. The statistical stability of these distributions was confirmed using linear mixed-effects models (LMM). For absolute power measures, the model revealed a highly significant interaction between cortical regions and clusters (P < 0.001). In contrast, spatial interactions for changes in relative power (Δδ and Δγ) were either not significant or only marginally significant (P = 0.413 and P = 0.063, respectively; Figs. 3E–H). Together, this distinct spectral segregation and geometric localization indicate that state-dependent yawning reflects more than a nonspecific physiological response activity. Instead, as the brain network transitions between states of consciousness, yawning appears to mark a specific and reproducible pattern of neurodynamic reconfiguration.

### Decoding Pre-Yawning Microstates Based on Nonlinear Complexity and Instantaneous Frequency Volatility

To further investigate the microscopic neuronal preconditions of yawning, we focused our analysis on a 30-second window (−30 to 0 s) preceding movement onset. Cluster-specific dynamic trajectories before yawning were identified by time-resolved Lempel-Ziv complexity (LZC) analysis (Fig. 4A). The slope of LZC evolution during the pre-yawning interval was used to stratify events, preventing opposing macroscopic dynamics from canceling each other out during cross-event averaging. Using this strategy, two distinct macroscopic dynamic modes were identified (Fig. 4B). Group 0 (n=59) exhibited a progressive decrease in LZC, indicative of network inhibition, whereas Group 1 (n = 37) showed a sustained LZC increase, consistent with a network rebound.

**Figure 4.**
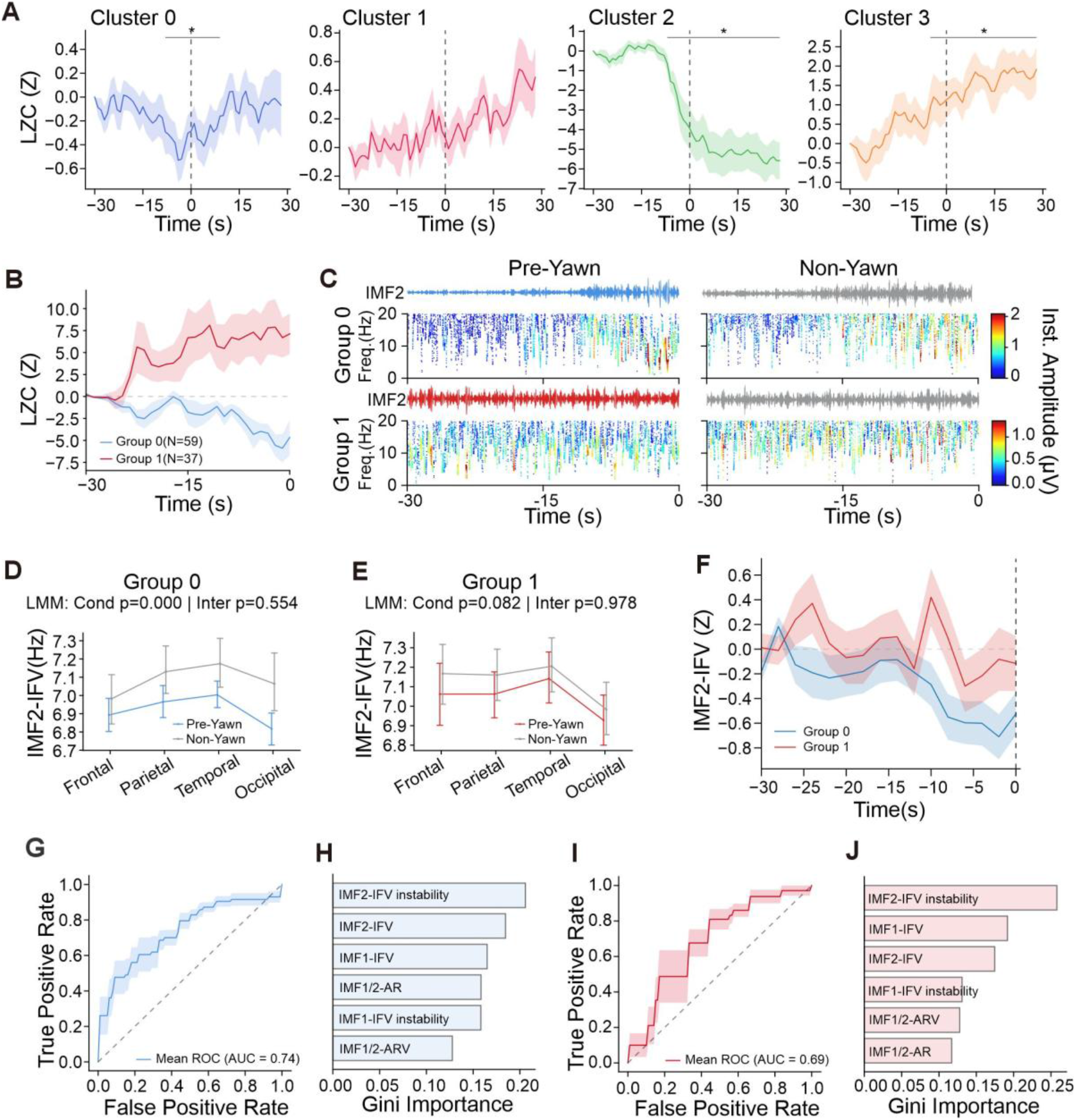
Pre-yawn microstate dynamics based on nonlinear complexity and instantaneous frequency volatility, and machine learning decoding. (A) Evolution of Lempel-Ziv complexity (LZC) for the four clusters around yawn onset (t = 0). The solid lines and shaded areas show the mean ± SEM for the events. The black lines with stars show the time windows where the data is significantly different from the baseline (−30 to −28 s) (cluster permutation test, P < 0.05). (B) Different types of LZC paths separated by their slope: Group 0 (complexity decreases, n = 59) and Group 1 (complexity increases, n = 37). (C) Examples of the Hilbert-Huang Transform (HHT) analysis for the states before a yawn and the control group. The top part shows the signal wave of the second intrinsic mode function (IMF2). The bottom part shows the Hilbert spectrum; the color bar indicates the instantaneous amplitude. (D, E) Mean values of the IMF2 instantaneous frequency volatility (IMF2-IFV) ± 95% CI for Group 0 (D) and Group 1 (E) in four brain areas. P-values show the main effect of state and the interaction effect calculated by the LMM. (F) Time tracks showing the IMF2-IFV (mean ± SEM) for Group 0 and Group 1 during the 30 seconds before the yawn starts. (G, I) ROC curves (Receiver Operating Characteristic) for the Random Forest models that decode the pre-yawn state for Group 0 (G) and Group 1 (I). The light shading represents the SEM between different folds. (H, J) Ranking of the Gini importance for the six main features used in the decoding models mentioned above.

To dissociate microstate-specific dynamics from macroscopic state drive, non-yawning control periods were selected by minimizing Euclidean distance within an 80-dimensional power spectral density space, thereby matching macro-level energy topologies for each yawning event. Using this rigorously matched baseline, EMD-HHT spectra showed pronounced limitations in pre-yawning microstate frequency switching (Fig. 4C). Subsequent quantitative analysis and continuous time tracking confirmed a dynamic decoupling between microscopic network flexibility and macroscopic state evolution (Figs. 4D–F). For statistical inference, linear mixed-effects models (LMM) with random intercepts for individual animals were applied. Relative to matched baseline periods, Group 0 showed a highly significant, global reduction in instantaneous frequency volatility of the second intrinsic mode function (IMF2-IFV), a measure reflecting intrinsic local network flexibility (P < 0.001). In Group 1, which exhibited a macroscopic rebound in state trajectory, microscopic network flexibility also failed to show robust recovery (P = 0.082; Figs. 4D, E). Continuous-time trajectories visibly confirm these statistical findings (Fig. 4F). Across both groups, the microscopic network consistently entered a brief period of flexibility inhibition, or “kinetic stagnation,” immediately prior to yawning onset, regardless of the direction of cortical arousal change. This result identifies a universal micro-dynamic signature preceding yawning.

Pre-yawning microstates were reliably distinguished from matched non-yawning control periods using a Random Forest classifier. Classification performance reached an Area Under the Curve (AUC) of 0.74 for Group 0 (accuracy = 64.66%; permutation test P = 0.0002; Fig. 4G) and 0.69 for Group 1 (accuracy = 66.22%; P = 0.0250; Fig. 4I). Feature importance analysis identified cortical IMF2-IFV and its second-order instability as the most discriminative predictors in both models (Figs. 4H, J). However, an important question remains. Reduced network flexibility also occurs during other rapid brain state transitions, such as anesthesia induction, yet yawning emerges only at specific, infrequent moments. This observation suggests that inhibition of intrinsic flexibility alone is insufficient to explain the selectivity of pre-yawning microstates. Instead, this narrowly defined temporal window likely reflects the interaction of additional neurodynamic parameters that gate the emergence of yawning.

### Cross-Species Spatiotemporal Decoupling of Micro-Complexity and γ-Band Power Preceding Anesthesia-Induced Yawning

To identify additional underlying dynamics beyond micro-flexibility inhibition, we focused specifically on the anesthetic induction period. This brief transition provides an ideal analytical window: its quick and highly structured progression from wakefulness to loss of consciousness enables the selection of baseline control segments that can be precisely matched to yawning periods. Because yawning is evolutionarily conserved, we expanded our analysis from the beagle model (n = 10 yawn/control pairs) to an independent human validation cohort (n = 9 pairs) to assess the cross-species universality of these neurodynamic precursors.

Within a 30-second window (−30 to 0 s) preceding yawning onset, we simultaneously extracted local γ-band power and Lempel-Ziv complexity (LZC). To generate a strict non-yawning control baseline, all trials containing induction-phase yawns were excluded from the 36 individual animal tests. We then identified the point of maximal LZC decline within the remaining induction-phase data; in control trials, this midpoint was identified as the pseudo-kinetic onset (t = 0). During anesthetic induction, this approach closely matched the baseline trajectory of macroscopic state collapse. By applying this stringent control framework, we were able to isolate yawning-specific micro-dynamics from the general effects of anesthetic induction. Together, this setup provides a high-fidelity platform for detecting transient decoupling between complexity and power across species.

Time-resolved trajectory tracking revealed distinct divergent evolutionary patterns under this precisely matched analytical framework (Figs. 5A, E and I, M), specifically between local high-frequency energy and global network complexity. Both yawning and control groups exhibited progressive LZC decay from −30 to 0 s, consistent with a general reduction in macroscopic arousal. However, pre-yawning microstates exhibited a brief increase in local γ-band power. To isolate baseline effects of passive state drift and quantify local energy mobilization, we computed the first time derivative (d/dt) of each signal (Figs. 5B, F). In contrast to the control group’s smooth, constant-velocity collapse, pre-yawning microstates displayed a nonlinear, accelerated deviation. We further applied Dynamic Time Warping (DTW) to align signal trajectories and compare their shapes. This result shows that the phenomenon is a genuine structural divergence in network dynamics rather than a simple temporal shift (Figs. 5C, G).

**Figure 5.**
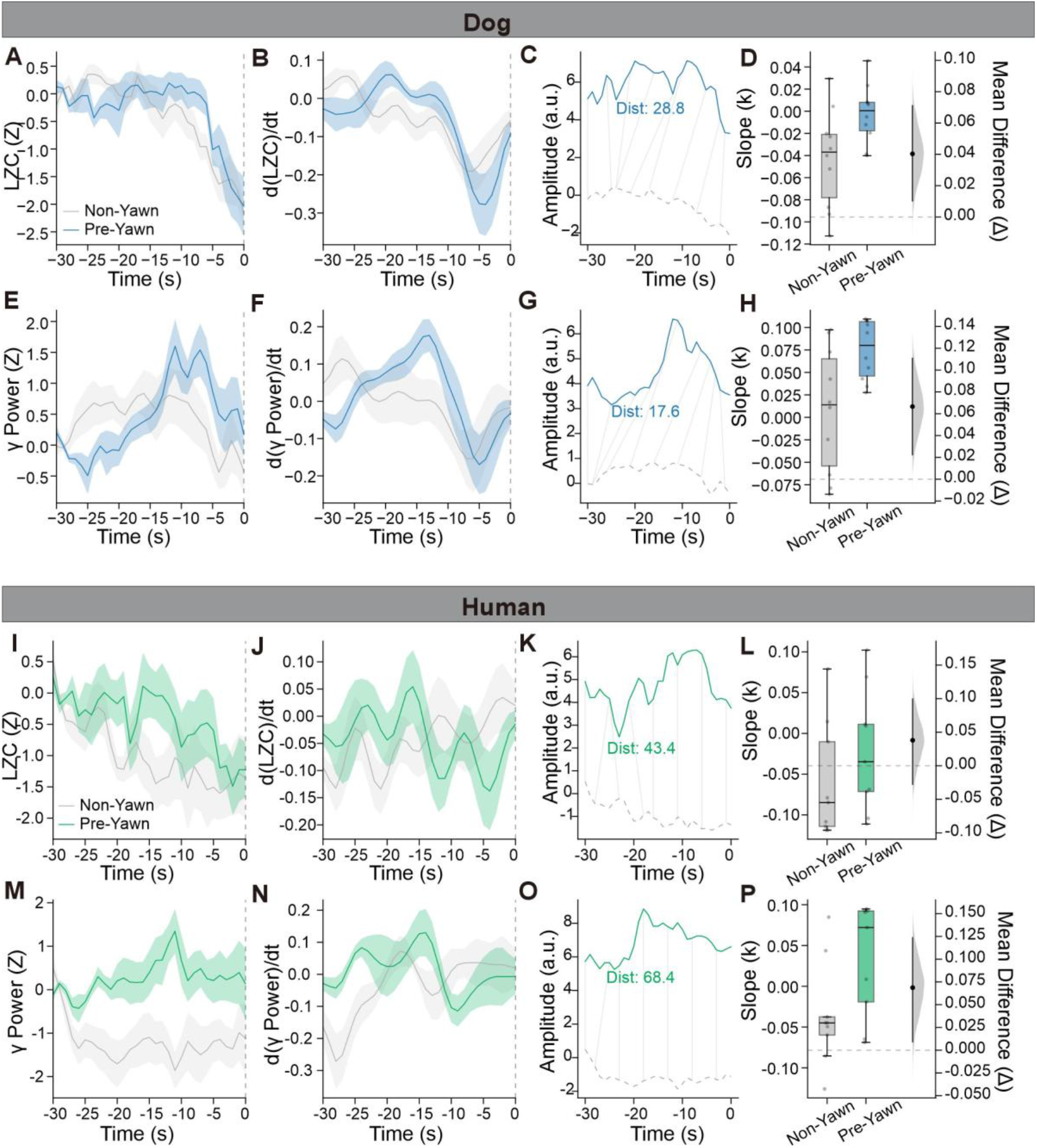
Cross-species dynamics and shape alignment of pre-yawn LZC and γ power during propofol induction. (A, E) Time tracks (mean ± SEM) showing LZC (A) and γ power (E) during the 30 seconds before a yawn starts. We used data from the beagle group (n = 10 yawn/control pairs). All values were normalized using Z-scores based on the baseline window (−30 to −28 s). (B, F) These show the first-order time derivatives (the rate of change) for the LZC and γ power data described above. (C, G) An example of matching the yawn signal (blue line) and the control signal (grey dashed line) using Dynamic Time Warping (DTW). The light grey lines show the best matching path between the two signals. The text inside the plot shows the calculated DTW distance. (D, H) Plots showing the slope (k) for the linear trend of LZC (D) and γ power (H). We calculated the slope for the time window between −30 s and −2 s. Black dots show the average difference between the yawn group and the control group. The vertical black lines show the 95% CI. (I–P) Results for the human group (n = 9 yawn/control pairs). The data processing and the statistical tests are exactly the same as the methods used in panels (A–H).

To further quantify temporal and spatial separation between macro and micro state dynamics, we calculated linear trend slopes (k) within a fixed time window (−30 to −2 s), reducing interference from movement. In the beagle model, pre-yawning γ-band power slopes were significantly higher than the control group (Δ = 0.0403, 95% CI: [0.0096, 0.0713]; Fig. 5H), as determined by bootstrap testing. LZC decay slopes also differed between conditions (Δ = 0.0666, 95% CI: [0.0213, 0.1124]; Fig. 5D). These important features were closely replicated in the human group.

This localized energy surge amidst global complexity reduction reflects a cortical “control effort” to preserve information integration. We term this phenomenon “Energy-Dynamics Mismatch” (EDM), defined as a marked disconnection between energy use and network flexibility. This mismatch captures the intrinsic tension opposing “Neural Inertia.” Yawning frequently emerges during periods when this underlying conflict intensifies.

### Neurodynamic Features and Cortical Decoupling of Pre-Yawn Microstates during Wakefulness

We expanded our investigation to yawning during natural wakefulness to determine whether similar network mismatches emerge during normal physiological behavior. To exclude residual anesthetic effects, EEG recordings were initiated seven days after the last propofol dose (Fig. 6A). During continuous behavioral monitoring, we collected 100 instances of spontaneous yawning. Analysis revealed a significant increase in high-frequency cortical activity preceding yawning onset. Time-resolved examination showed that γ-band (30–45 Hz) power remained elevated above baseline prior to yawning, peaking at approximately −10 s before gradually declining as the yawn itself approached (Fig. 6B). We also calculated total γ energy over the 30-second pre-yawn interval, which demonstrated a highly significant increase. Compared to the precisely matched non-yawning control periods, this change was significantly different (Fig. 6C, P < 0.001).

**Figure 6.**
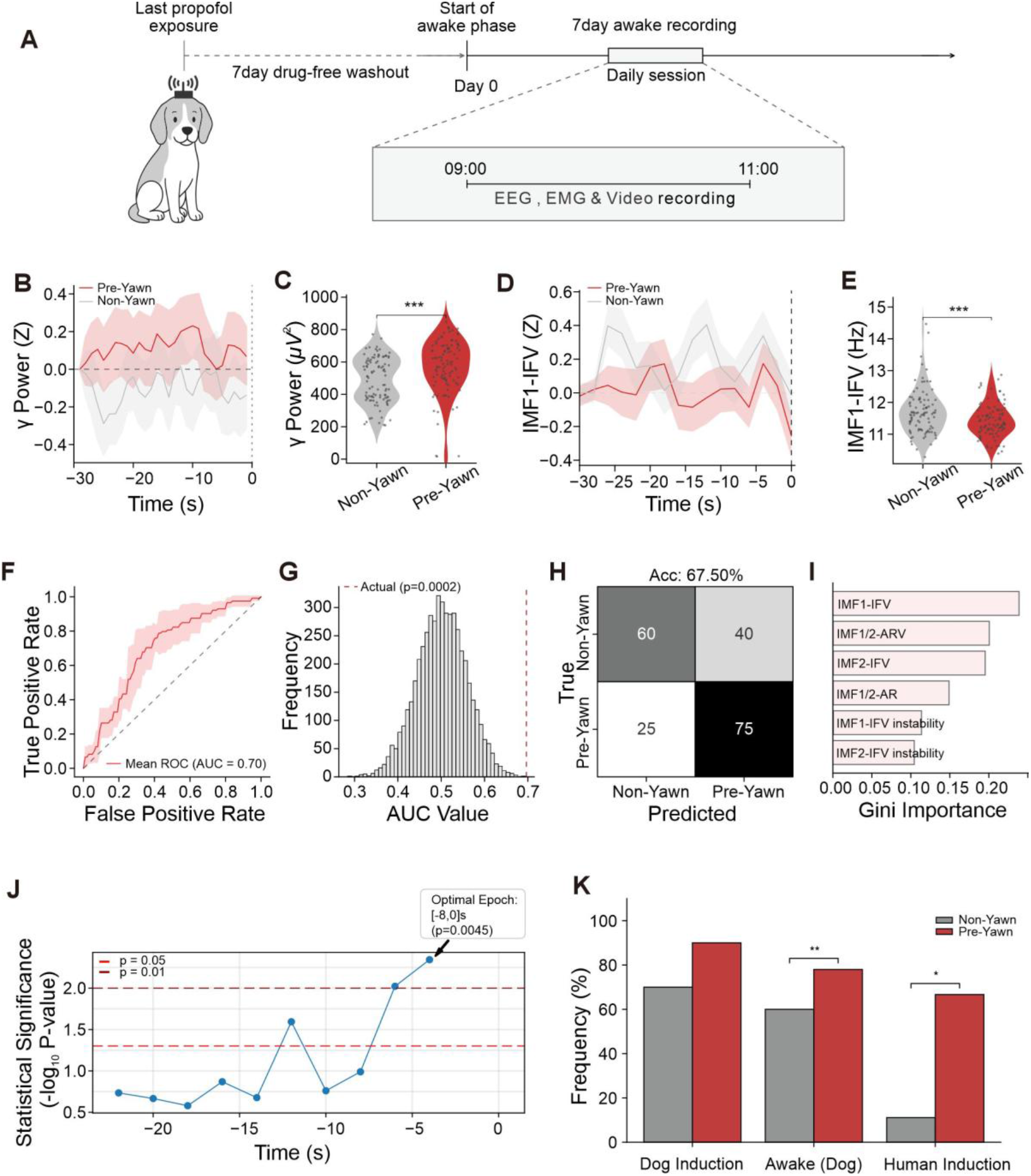
Decoupling and decoding of pre-yawn microstates during wakefulness. (A) Schematic diagram of the longitudinal waking period experimental design. (B, C) γ power tracks before a yawn compared to the control period (B) and the comparison of γ energy (C) when awake. All data were normalized to Z-scores using the baseline window (−30 to −28 s). The plots show the mean ± 95% CI. ***P < 0.001. (D, E) Changes in IMF1-IFV while awake (D) and the statistical comparison (E). We used the same baseline method as in panel B. The data show the mean ± SEM. ***P < 0.001. (F–I) Using a Random Forest model to find the state before a yawn. This part includes the average ROC curve (F), the distribution of 5,000 permutation tests (G), the confusion matrix (H), and the ranking of feature importance (I). (J, K) Using the data to find the best time window for detecting Energy-Dynamics Mismatch (EDM) (J) and a comparison of EDM detection rates in different species (K). *P < 0.05, **P < 0.01, based on Fisher’s exact test.

In contrast, the instantaneous frequency volatility (IFV) decreased concurrently with rising high-frequency power. This separation of dynamics is clearly illustrated in the tracking plots (Fig. 6D). Before a yawn, IFV of the first intrinsic mode function (IMF1-IFV) remained consistently lower than baseline and lower than the more flexible oscillatory dynamics observed during control periods. As yawning onset approached, IMF1-IFV dropped very quickly. Quantitative analysis confirmed a highly significant overall reduction in IMF1-IFV across the entire pre-yawning window (Fig. 6E, P < 0.001), reflecting a marked decrease in intrinsic local network flexibility.

Applying the identical Random Forest classification framework used for the anesthesia data, we successfully distinguished pre-yawning microstates from matched controls during wakefulness, achieving a mean AUC of 0.70 (permutation test P = 0.0002; Figs. 6F, G) and an overall accuracy of 67.50% (Fig. 6H). Feature importance analysis identified IMF1-IFV as the strongest predictive variable (Fig. 6I).

To quantify energy-dynamics decoupling at the single trial level, we implemented a data-driven sliding window approach (8-s window, 2-s step). This analysis revealed that the Energy-Dynamics Mismatch (EDM) was most pronounced in the 8 seconds immediately before the yawn started (Fig. 6J), which we selected as the focal interval for subsequent analyses. Within this window, EDM events were identified for each individual using two thresholds: γ power exceeding the 75th percentile of baseline values and IFV falling below the 25th percentile. Cross-species comparisons confirmed the robustness of this effect (Fig. 6K). In both awake dogs and humans undergoing propofol induction, EDM events occurred significantly more frequently prior to yawning than during matched control periods. Together, these findings demonstrate that pre-yawning dynamics during wakefulness are distinct from general state changes and that the EDM effect is tightly confined to the temporal vicinity of yawning onset.

## Discussion

We investigated the spatiotemporal dynamics of yawning across different arousal states. Our results show that yawning is not a simple, uniform brain activity (20). Instead, it consistently coincides with a specific micro-dynamic decoupling process in the brain. In this process, local cortical energy increases rapidly (e.g., high-frequency γ activity), while network flexibility decreases. We quantify this flexibility using instantaneous frequency volatility (IFV) and define this transient mismatch between elevated local energy demand and delayed microstate switching as “Energy-Dynamics Mismatch (EDM).”

Propofol-induced yawning exhibited marked asymmetry on a macro-temporal scale. Specifically, 89.2% of yawning events clustered during the recovery phase. In pharmacology, the difference in drug levels required to induce versus regain consciousness is referred to as “neural inertia” (21, 22). During recovery, residual anesthetic effects compete with endogenous arousal drive, prolonging the transition window (23). The clustered yawning events observed during this phase likely reflect periods of intense macroscopic reorganization as the system overcomes this neural inertia.

Our micro-dynamic neural features, extracted using EMD-HHT, provide quantitative insight into these macro-state conflicts. Traditional EEG studies often associate yawning with increases in slow wave activity and suppression in the α-rhythm (24). In contrast, EMD-HHT captures quick, transient changes by breaking down signals into intrinsic modes and estimating instantaneous frequency (25, 26). Using this approach, we revealed previously hidden pre-yawning dynamics that remain consistent with established signatures of low arousal (27). Specifically, we observed rapid increases in γ power and lower flexibility (lower IMF-IFV) in the brain just before a yawn starts. Previous studies linked γ-band activity and high energy needs of local neurons (28, 29). The simultaneous increase in γ power and the reduction in flexibility may therefore reflect elevated energetic requirements for information processing (30, 31), consistent with the theoretical concept of “control effort.” Importantly, time-resolved trajectories (Fig. 5 and 6) demonstrate that these high-frequency changes are not EMG artifacts. Cortical γ power consistently declined before the actual physical movement began, confirming that the elevated energy precedes motor execution.

We further observed that the intrinsic modes (IMFs) carrying transient flexibility suppression exhibited a state-dependent frequency shift. Specifically, this “decoupling” moves to the highest frequency mode (IMF1) when the animal is awake, but during anesthesia induction and recovery, the second high-frequency mode (IMF2) is more prominent. Interestingly, low-frequency waves have been shown to become the main part of brain activity when high-frequency activity is suppressed (32, 33). On the other hand, the brain naturally has more activity and higher-frequency waves when it is very alert and awake (34, 35). So, this clear shift in frequency bands shows how the network always adjusts its main wave mode under different levels of wakefulness.

Together, these findings provide a new framework for understanding yawning as a conserved, cross-species behavior. In contrast to traditional theories that frame yawning as a passive byproduct of sleepiness or thermoregulation (36), our data indicate that the pre-yawning brain state reflects an active conflict during large-scale state reorganization. Although yawning is classically associated with brainstem and hypothalamic structures such as the paraventricular nucleus (PVN) (37, 38), transitions between arousal states involve whole-brain network dynamics (39, 40). The destabilization and reorganization of cortical dynamics frequently precede or accompany the release of subcortical primal circuits (41, 42). As local cortical regions exhibit increased energetic demand during periods of reduced global control (43), dynamic decoupling emerges across hierarchical levels. Thus, yawning reflects a specific behavioral manifestation of constrained network flexibility rather than a simple reflex.

Finding these hidden brain patterns was enabled by the pharmacological model and the use of rigorously matched baselines. Anesthetic manipulation amplified resistance during state transitions, facilitating detection of otherwise subtle dynamics (44, 45). Importantly, EDM was replicated in both dogs and an independent human cohort, supporting its potential role as a conserved mammalian signature of state transition dynamics (46). Clinically, conventional EEG‐based monitoring tools that rely on slow‐wave metrics often detect state transitions only after substantial reorganization has occurred(47). Our findings suggest that increases in local energy demand coupled with abrupt reductions in network flexibility (IMF‐IFV) may precede macroscopic EEG changes, offering a potential early indicator of network instability.

Despite the strong temporal association between microdynamics and yawning, several limitations warrant consideration. First, as an observational study, we cannot establish a causal link between EDM and interactions between cortical and subcortical yawning circuits. Second, generalizability across anesthetic classes remains to be tested. Future investigations should integrate high-resolution fMRI (48) or multi-region microelectrode recordings in (49) with circuit-specific perturbation methods (50) to clarify causal mechanisms linking EDM to yawning centers.

In summary, the neural signatures identified here provide new quantitative insight into how complex brain networks reorganize during instability and state changes and suggests a foundation for future development of early warning systems for BCI based on brain state changes in the future.

## Materials and Methods

### Experimental Model and Data Acquisition

We performed longitudinal recordings of epidural EEG and intramuscular masseter EMG in six beagle dogs during propofol-induced state transitions. Yawning behavior was objectively identified using markerless kinematic tracking (DeepLabCut). To assess cross-species universality, an independent clinical EEG dataset comprising nine matched yawn/control segments was analyzed. All animal and human procedures were approved by institutional ethics committees (see SI Appendix).

### Feature Extraction and Predictive Modeling

Macroscopic neural trajectories were reconstructed using principal component analysis and generalized additive models. To resolve millisecond-scale microdynamics beyond linear time–frequency limits, we applied empirical mode decomposition (EMD) with Hilbert spectral analysis. Network flexibility was quantified via instantaneous frequency volatility (IFV). Pre-yawn microstates were classified using a Random Forest algorithm with cross-validated feature importance.

### Detailed Methods

Comprehensive protocols covering surgical workflows, signal preprocessing, clustering pipelines, Energy–Dynamics Mismatch (EDM) quantification, and statistical frameworks are provided in SI Appendix, Materials and Methods.

## Acknowledgments

This work was supported by the Wu Jieping Medical Foundation (No. 320.6750.2024‐05‐14, Wenfei Tan); by the National Natural Science Foundation of China (No. 82171187, Wenfei Tan); and by the China zhongguancun Precision Medicine science and technology foundation (No. 2024-11--085 to L.Q.).

