## Supporting Information for "Yawning reveals energy-dynamics mismatch and neural inertia across state transitions"

Xiangshang Meng[a], Qi Sun[a], Muzi Li[a], Xuanze Li[a], Guangtai Liang[a], Ling Qin[b]* and Wenfei Tan[a]*
[a] Department of Anesthesiology, the First Hospital of China Medical University, Shenyang 110001, China
[b] School of Life Sciences, China Medical University, Shenyang 110122, China
*Ling Qin and Wenfei Tan

**This PDF file includes:**

Supporting text
Figures S1

### Supporting Information Text

### Extended Results and Methodological Validation

**Exploratory Time-Frequency Analysis and EMD-HHT Validation.** To initially assess microscopic dynamics associated with anesthesia-related yawning, we applied Morlet wavelet transforms for conventional time-frequency analysis of single-trial EEG signals (Fig. S1C). Exploratory inspection revealed distinct trends between the two groups. Early in the pre-yawning window, cortical energy in Group 0 alternated between high and low frequency bands while the macroscopic network exhibited inhibitory dynamics. However, near yawning onset (about −8 s), the flexibility of frequency switching became severely limited, as evidenced by a continuous shift of energy toward lower frequencies. In contrast, Group 1 showed consistently low local frequency variability throughout the window. This dimension appears critical for characterizing microstate dynamics preceding yawning.

To more precisely quantify this behavior, we employed EMD-HHT, which is well suited for analyzing nonstationary brain signals that challenge traditional linear approaches. To ensure feature consistency, we examined the center frequencies of Intrinsic Mode Functions (IMFs) derived from frontal channels across multiple trials. The results demonstrated a robust "band-anchoring effect". Despite substantial variability in background consciousness states across the 96 yawning events. Specifically, a clear separation was observed between the low-frequency mode (IMF2) and high-frequency mode (IMF1), anchored near the θ/α band (7.1 ± 2.1 Hz) and the γ band (33.6 ± 2.8 Hz), respectively. This stability was essential for ensuring the reliability of subsequent analyses. Moreover, single trials from Groups 0 and 1 exhibited highly similar frequency distributions (Fig. S1F). Consistent with anesthesia results, EMD analysis during wakefulness revealed a clear and complete separation of frequency components (Fig. S1G). IMF1 captured high-frequency activity and remained anchored within the γ band (36.9 ± 2.2 Hz), while IMF2, the low-frequency mode, was fixed at 6.5 ± 1.9 Hz. Because these frequency bands remained stable across different background states, the common issue of "mode mixing" in EMD was effectively avoided.

**Cross-Band Synergy and Spatial Specificity.** When combined with parallel component analysis, this state-dependent micro-flexibility reconfiguration revealed strong cross-band synergy (Figs. S1D, E). In Group 0, IMF1 and IMF2, representing high- and low-frequency microstates, respectively, both showed a highly significant global attenuation in flexibility ($P < 0.001$ for both state main effects). In contrast, Group 1 exhibited blunted multi-scale dynamics with no obvious fluctuation (main effects of condition for IMF1 and IMF2: $P = 0.699$ and $P = 0.082$, respectively). Notably, neither group showed any significant "state × region" interaction effects (P-value range: 0.491 to 0.978), indicating that pre-yawning flexibility reconfiguration is not spatially localized. Instead, it reflects a brain-wide dynamic process that depends strongly on the underlying macroscopic arousal state.

**Machine Learning Feature Selection.** Given the observed brain-wide consistency, we simplified the input features for our single-trial machine learning classification to identify key predictive parameters. Features were strictly restricted to frontal channels to reduce collinearity inherent in high-dimensional spatial representations. Furthermore, a −32 to −2 s time window was selected to strictly avoid interference from motor-ready potentials and early EMG artifacts prior to yawning onset. Using four frontal channels, we constructed a six-dimensional feature space encompassing IMF-IFV and its instability metrics to train the Random Forest classifier.

### Materials and Methods

**Experimental animals.** The study used six healthy female Beagle dogs (6–12 months old, weight 8–10 kg). The animals were kept in standard rooms and had free access to food and water. The room temperature was 22 ± 2°C, the humidity was 50%–60%, and a 12-h light/dark cycle was used.

The Animal Experiment Ethics Committee of China Medical University approved all the procedures (Approval No. 20251142) and this work followed the Guide for the Care and Use of Laboratory Animals.

**Human EEG Dataset.** We used human EEG data from a database at the First Hospital of China Medical University. The Medical Ethics Committee approved the data collection plan (Approval No. [2025]2025-507-2), and the study was registered with the Chinese Clinical Trial Registry (ChiCTR2500103843). All participants provided written informed consent before data collection began. This included nine yawning events and non-yawning segments that were carefully matched. The participants were between 18 and 65 years old with American Society of Anesthesiologists (ASA) physical status I or II. The exclusion criteria included any history of neurological or psychiatric disorders.

**Stereotaxic Landmark Implantation and MRI Targeting.** Before the electrode surgery, we implanted markers to have a fixed reference for the MRI. We anesthetized the animals with propofol (8 mg/kg, i.v.), shaved the scalp, cleaned it with chlorhexidine, and made an opening along the midline. We separated the skin and tissue to find Bregma, then put a small titanium screw (non-ferromagnetic) at the Bregma to use as a reference point before closing the scalp in layers. The animals recovered for three days with routine antibiotics and medicine for pain. After they recovered, the animals had an MRI scan (Siemens Prisma 3.0T) while under propofol anesthesia. We took high-resolution 3D T1 images (3D MPRAGE; 0.6 mm slice thickness; $0.8 \times 0.8$ $mm^2$ pixel spacing), and uploaded the DICOM data into 3D Slicer software. The metal marks from the titanium screws showed us the starting point (origin) in the images. Using a dog brain atlas, we chose 16 target sites in the frontal, parietal, temporal, and occipital areas and made a specific drilling map for each animal. The Bregma was the center coordinate (0, 0), while the surface coordinates for the 16 brain sites (AP, ML; mm) were: bilateral frontal (AP +20.0, ML ±8.0; AP +10.0, ML ±12.0), bilateral parietal (AP −10.0, ML ±10.0; AP −20.0, ML ±12.0), bilateral temporal (AP −15.0, ML ±20.0; AP −25.0, ML ±22.0), and bilateral occipital (AP −32.0, ML ±8.0; AP −40.0, ML ±10.0).

**Electrode Array Assembly and Implantation.** The integrated recording array was assembled before surgery. It included 16 insulated silver wires (cortical EEG, A-M Systems, #785500; 1–2 mm stripped), 2 insulated nichrome wires (masseter EMG, #793200; 3–5 mm stripped), and 2 silver wires for reference and ground (insulation stripped over a large area). These were soldered to a multi-channel connector and sterilized. Animals received propofol (8 mg/kg, i.v.) for induction and isoflurane (1.5%–2.0%) for maintenance. The head was fixed in a stereotaxic frame. We exposed the skull and drilled 16 small bone windows using MRI coordinates. The dura mater remained intact. EEG electrodes were placed in the epidural space and sealed with bone wax. EMG wires were tunneled subcutaneously to the cheek, and the exposed ends were inserted into the masseter muscle belly using 4-0 polypropylene sutures. The poles were spaced 2 mm apart for bipolar recording and 4–5 titanium screws were implanted for reference and ground. Dental acrylic secured the wires and connectors into a head cap. Incisions were sutured with 4-0 nylon. Animals received antibiotics and analgesics for three days.

**Experimental Design and Drug Administration.** We used a longitudinal repeated measures design to establish a standardized model of anesthesia state transitions. In a pilot study (n = 3), we compared 8 mg/kg and 4 mg/kg propofol using a randomized crossover design. Propofol was infused intravenously at a constant rate over 1 min via a KL-702 dual-channel syringe pump (Beijing Kelijianyuan Medical Technology Co., Ltd., China). A 3-day washout period separated each dose. For the main experiment (n = 6), we selected the 4 mg/kg dose. Each animal finished three testing cycles, giving us 36 anesthesia records and 18 awakening records. Every trial followed a standard procedure: 4-h fasting, recording baseline EEG, 1-min propofol infusion, and simultaneous marking of important behavioral points, such as loss of response (LOR), opening eyes, and walking. To avoid drug buildup and to look at network mismatch features in the awake state, we waited for 7 days after the last propofol dose. After this, we monitored the baseline activity every day for 7 days in a row during a fixed time window (09:00–11:00).

**Simultaneous EEG, EMG, and Video Recording.** The brain and behavior data was collected in a quiet room that was protected from electrical interference. Before starting, we connected the Hermes wireless transmitter (Bio-Signal Technologies, USA) to the electrodes on the dog's head. The recording began after the animals stopped moving and their physical states were stable. We recorded the brain (EEG) and muscle (EMG) signals continuously at 1000 Hz. Video was taken with a high-definition camera at 30 fps and 1080p resolution using OBS Studio software. To make sure everything was perfectly synchronized, a special AutoHotkey script started both the signals and the video at the same time. The animals stayed within 2 m of the wireless receiver to keep the signal stable during the experiment.

**Yawning Behavior Identification and Kinematic Analysis.** We analyzed yawning behavior from the video source using DeepLabCut (v2.3.5) to track movements without markers. We collected the 2D coordinates of the jaw, mouth corners, and the tip of the nose to see how the face moved. To determine yawns from other mouth movements like chewing or licking, we used strict rules: the jaw opening angle (θ) was fast and nonlinear, and each mouth-opening event had to last longer than 1 s. We ensured that the onset of mouth opening coincided with the onset of masseter EMG bursts. We set the start of these EMG bursts as time zero ($t = 0$) for every yawn. This time point served as the anchor for extracting time-series data for subsequent analysis.

**EEG and EMG Data Preprocessing.** MNE-Python (v1.10.1) software was used to process the EEG and EMG data. First, a channel-location file containing the Cartesian coordinates (x, y, z) for each electrode was generated and linked to the corresponding data channels. We filtered the data (0.5–80 Hz) and changed the sampling rate to 250 Hz. To remove noise that was not from the brain, we used Independent Component Analysis (ICA) to carefully look at the time plots, power graphs, and scalp maps by eye to find and remove these noise parts. Components related to eye movements, heartbeats, or head muscle noise were removed. Then, we re-referenced the cleaned EEG data to the common average (CAR). For masseter EMG signals, raw data were first band-pass filtered (20–400 Hz, fourth-order zero-phase Butterworth) to retain motor unit action potentials while suppressing movement artifacts and high-frequency noise. The filtered signals underwent full-wave rectification to preserve the amplitude modulation of muscle activity. A linear envelope was then extracted by applying a 10 Hz low-pass filter (fourth-order zero-phase Butterworth) to the rectified signal. This envelope shows the strength of the masseter muscle activity.

**Neural State Space Construction and Trajectory Analysis.** To measure brain state dynamics during anesthesia induction and recovery, we calculated the power spectral density (PSD) of segmented EEG signals using Welch's method in SciPy (v1.14.1). We used a 5.0-s sliding window with a 2.0-s step to achieve the relative power for five frequency bands: δ (0.5–4 Hz), θ (4–8 Hz), α (8–13 Hz), β (13–30 Hz), and γ (30–45 Hz). We put the spectral features from all channels together and normalized them using Z-scores. Then, we used Principal Component Analysis (PCA) in Scikit-learn (v1.7.2) to reduce the data to three dimensions. This formed the neural state space. Then, using a Savitzky-Golay filter (length 51, order 3) we smoothed the 3D paths to reduce noise. We measured how the state changed by calculating the Euclidean distance between the current brain state and the center of the anesthesia state. This center was considered the average state when the animal lost response (LOR). Finally, we used a Generalized Additive Model (GAM) in pyGAM (v0.12.0) with 25 spline functions to fit the distance data. The confidence intervals came from this model fit.

**Yawning-Related Neural Dynamics Clustering Analysis.** Continuous EEG data was extracted from −30 s to +30 s around each yawning onset. We used MNE-Python to do time-frequency analysis with Morlet wavelets. The frequencies were from 1 to 45 Hz with 45 points, and we set the wavelet cycles to f/2 for each frequency. To characterize network state changes around yawning, we built a 64-dimensional feature vector for each yawning event by combining 4 core metrics from all 16 EEG channels. For every channel, these four specific measures included the change in relative δ power before versus after the yawn onset ($\Delta Power\_\delta$); the change in relative γ power before versus after the yawn onset ($\Delta Power\_\gamma$); the average δ power calculated over the entire 60-s time window; and the average γ power calculated over the entire 60-s time window. We normalized all features using Z-scores to make them comparable. Then, we used K-Means clustering in Scikit-learn to find patterns in the data. We determined the optimal number of clusters (K = 4) using the elbow method based on within-cluster sum of squares (WCSS), supplemented by the silhouette coefficient. K-Means clustering was then performed with 10 random initializations (n_init = 10), and the solution with the lowest inertia was retained to ensure convergence stability. To check if the clusters were strong, we used UMAP from umap-learn (v0.5.9) to map the high-dimensional features into a 2D space with n_neighbors = 8 and min_dist = 0.3.

**Nonlinear Complexity and Micro-Dynamics Feature Extraction.** We used Lempel-Ziv complexity (LZC) to assess cortical network information integration. Signal median served as the binarization threshold and a 2.0-s sliding window with 1.0-s step was applied. Data were normalized using baseline periods from −30 s to −28 s before yawning onset. To overcome resolution limits of traditional linear time-frequency analysis, we performed empirical mode decomposition (EMD) on broadband signals (1.0–45.0 Hz) using PyEMD (v1.6.4). Standard EMD is prone to mode mixing when processing broadband non-stationary signals with intermittent noise. We applied strict pre-filtering and parameter constraints to ensure decomposition quality. Before EMD, EEG signals were bandpass filtered (0.5–45.0 Hz) to remove high-frequency muscle transients and ultra-low-frequency baseline drift. Maximum decomposition layers were limited to 4 (max_imf = 4), forcing the algorithm to extract dominant oscillation patterns within restricted frequency bands. This strategy reduced pseudo-mode generation while keeping computational efficiency for large-scale high temporal resolution sliding window scans, compared to expensive ensemble EMD variants. From the first two intrinsic mode functions (IMF1 and IMF2), we extracted analytic signals via Hilbert transform in SciPy. Instantaneous phase was unwrapped and differentiated to obtain instantaneous frequency (IF). Neural flexibility was defined as IF volatility (IFV), calculated as the standard deviation of instantaneous frequency over time. We also extracted first-order time derivatives of complexity and high-frequency energy using 1D Gaussian smoothing ($\sigma$ = 2.0) with discrete gradient operations. These derivatives measured instantaneous change rates (velocity) of network state evolution.

**Machine Learning Predictive Analysis.** A Random Forest classifier was built using Scikit-learn. The model had 300 trees, and we set the maximum depth to 5 to avoid overfitting. For input data, we used frontal EEG channels to calculate a six-dimensional feature set using sliding windows. Specifically, we calculated the instantaneous frequency volatility (IFV) and its instability over time for the first and second modes (IMF1 and IMF2). To look at the balance between high-frequency and low-frequency energy, we also included the amplitude ratio (IMF1/2-AR) and its variability (IMF1/2-ARV). To ensure the data was handled correctly and to avoid errors from similar samples, we used Stratified Group K-Fold Cross-Validation. The Area Under the Curve (AUC) measured how well the model performed. We also did 5000 random permutation tests to check for statistical significance. Finally, we used Gini importance to see which features were the most important in the model.

**Cross-Species Dynamics Feature Alignment and Statistical Analysis.** We compared how brain states change in different species using Dynamic Time Warping (DTW) with the fastdtw (v0.3.4) package to measure and match the differences in the paths between yawning events and control periods. To see how network measures changed before the yawn started, we looked at the data from −30 s to −2 s. This specific time window was used to avoid noise from movement just before the action. We calculated the trajectory slopes (k) using linear regression in SciPy.

**Single-Trial Energy-Dynamics Mismatch (EDM) Quantitative Analysis.** We smoothed the continuous EEG signals using a 1-s sliding average window. We took out the local $\gamma$ band power (30–45 Hz) and the Instantaneous Frequency Volatility (IFV) from the Intrinsic Mode Functions (IMF). The start of the yawn was the reference point (t = 0) and we used the time from −30 s to −28 s as the baseline for each subject. To find the best window for seeing changes in features, we looked through the data of the awake Beagle dog group. We used an 8-s sliding window with a 2-s step, starting from −26 s. From this, we set the main window as [−8.0 s, 0.0 s]. Then, we measured all single trials in the same way for all species. The main IMF bands were picked specifically for each experimental state. To classify the events, we used two threshold rules. For an EDM event to be positive, two things had to happen in the test window: first, the maximum γ power had to be higher than the 75th percentile of the baseline; second, the minimum IFV had to be lower than the 25th percentile of the baseline. Finally, we counted the EDM event frequencies and compared them between the different groups. We compared yawning events against control periods that were matched strictly.

**Statistical Analysis.** To determine whether yawning happened differently between the anesthesia stages (induction vs. recovery), we changed the raw yawn counts into rates per minute. This helped us fix the bias from different stage lengths. We used linear mixed-effects models (LME) to analyze these rates. The anesthesia stage was a fixed effect, and Animal ID was a random effect. This helped us manage repeated measurements from the same dogs. For data that did not follow a normal distribution, like the energy values, we used Kruskal-Wallis tests. Then, we compared pairs of groups using Mann-Whitney U tests from SciPy. All these tests were corrected using the Benjamini-Hochberg method (FDR-BH). For time-series data like the LZC paths, we used cluster-based permutation tests from MNE-Python. This was done to fix problems from having many tests and data being related over time. We ran 1000 random tests with a threshold of $P < 0.15$ to find the clusters. For count data, such as the EDM event frequency between yawning and control groups, we used one-sided Fisher's exact tests with the alternative hypothesis specified as directional (testing for a significantly higher frequency in the yawning group). We also calculated the average differences with 95% confidence intervals (95% CI) using 5000 bootstrap samples and made density plots from these samples. Unless we mention otherwise, we used a significance level of $P < 0.05$ for all tests.

### Figures

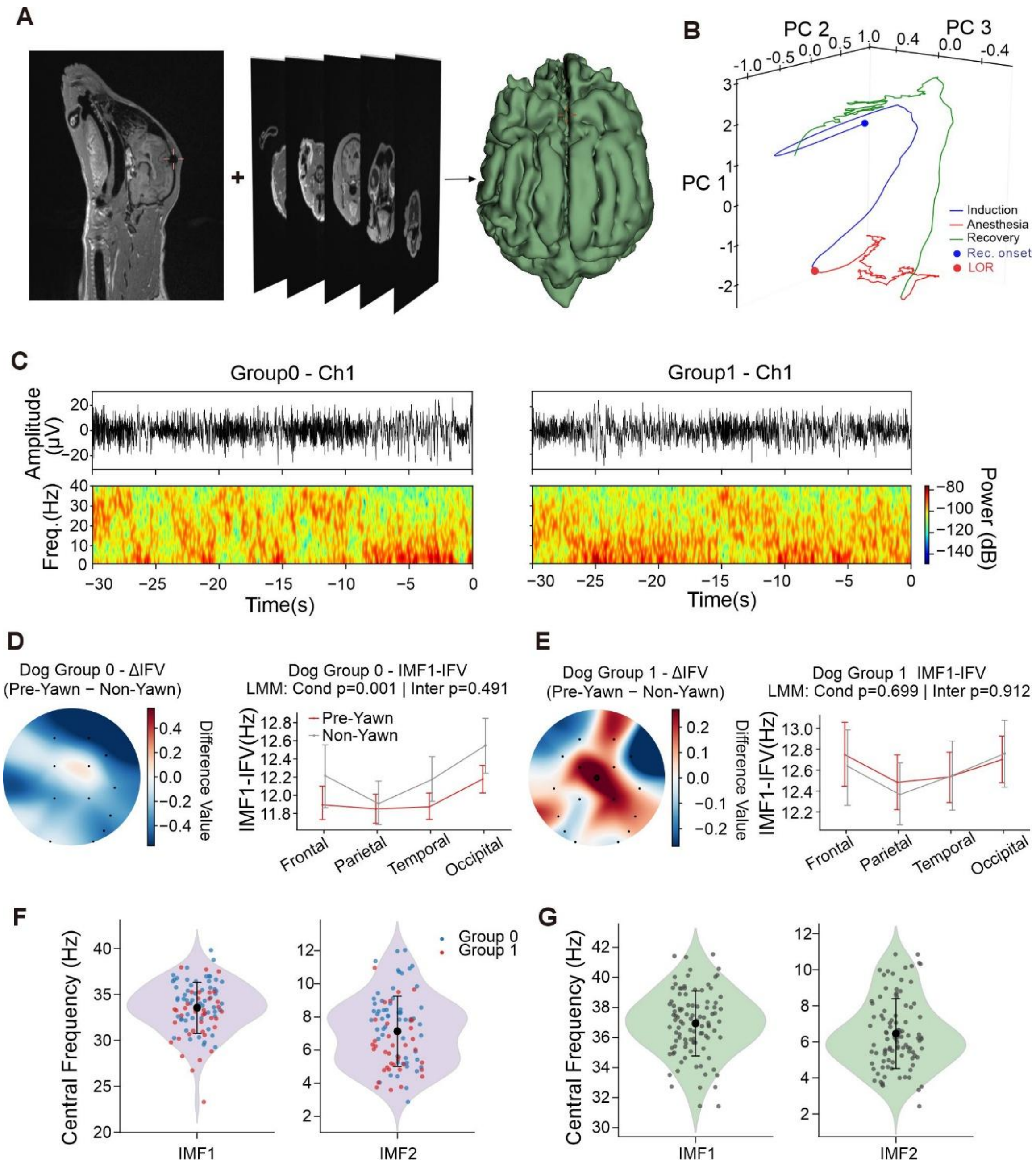


**Fig. S1. Finding brain locations, paths of brain states, and checking the frequency of EMD modes.** (A) Using MRI to find locations and build a 3D brain model of the beagle. From left to right: sagittal slices, coronal slices, and a 3D model of the whole brain surface. The red crosshair shows the physical starting point at Bregma. (B) 3D paths of brain states (PCA manifold) during anesthesia for one beagle. The blue, red, and green lines show the induction, maintenance, and recovery phases. Blue and red dots mark the start of record (Rec. onset) and the loss of response (LOR). (C) Time-frequency dynamics of representative single trials of Group 0 (left) and Group 1 (right) before yawning (−30 to 0 s). The continuous time-domain waveform of Channel 1 is shown above, and the time-domain spectrogram of the Morlet wavelet is shown below. (D, E) Data for Group 0 (D) and Group 1 (E). On the left: maps showing the difference in IMF1-IFV (ΔIFV) between the time before a yawn and the control period. On the right: average IMF1-IFV ± 95% CI for four brain areas.

P-values show the effects calculated using the LMM. (F, G) The frequency anchoring effect of EMD. These show the center frequencies for IMF1 and IMF2 during anesthesia (F) and when the animal is awake (G). We used the baseline window (−30 to −28 s) to find these features. In (F), blue and red dots represent Group 0 and Group 1.
